# Lateralised modulation of posterior alpha oscillations by closed loop auditory stimulation during memory retention

**DOI:** 10.1101/2025.05.07.652437

**Authors:** Henry Hebron, Radost Dimitrova, Valeria Jaramillo, Derk-Jan Dijk, Ines R. Violante

## Abstract

Alpha oscillations have been implicated in the maintenance of working memory representations. Notably, when memorised content is spatially lateralised, the power of posterior alpha activity exhibits corresponding lateralisation during the retention interval, consistent with the retinotopic organisation of the visual cortex. Beyond power, alpha frequency has also been linked to memory performance, with faster alpha rhythms associated with enhanced retention. These findings position alpha oscillations as a promising target for neuromodulation.

In this study, we demonstrate that although alpha frequency is not typically lateralised in a retinotopic manner during working memory retention, such lateralisation can be externally induced. Using alpha closed-loop auditory stimulation (αCLAS), and leveraging the phase-dependent responsiveness of alpha oscillations to sound, we successfully modulated alpha frequency asymmetrically between the visual cortices. The extent of induced frequency lateralisation was associated with the behavioural asymmetry in task performance.

## 1. Introduction

Working memory (WM) refers to the temporary retention and re-processing of information from the very recent past (Baddeley and Hitch 1974; Baddeley 2003), and functions as a substrate upon which we are able to concurrently process and synthesise disparate recalled ideas, rendering it fundamental to function in the world. WM declines with age (Balota et al. 2000; Zacks et al. 2000; Schneider-Garces et al. 2010; Auer et al. 2024), and WM deficits are a prominent feature of several brain-related diseases, such as Alzheimer’s disease (Becker 1988; BADDELEY et al. 1991).

Even in healthy individuals, WM has limited capacity (Luck and Vogel 1997; Cowan 2001; Baddeley 2003), which must be managed well if it is to be useful. One of the ways in which this can occur is by prioritising certain memories over others, an act that most of us carry out automatically, but can be considered as an explicit and conscious allocation of internal attention (Griffin and Nobre 2003).

A common paradigm used to explore this prioritisation/allocation in WM is cueing (Griffin and Nobre 2003; Ma et al. 2014; Souza and Oberauer 2016), in which a to-be-remembered visual sample (e.g., an assortment of coloured squares) is presented to a participant and they are instructed to prioritise remembering a particular portion of the sample (e.g., squares from the right-hand side). There is clear benefit of cueing on recall (Griffin and Nobre 2003; Ma et al. 2014), even if the cue is delivered after the to-be-remembered sample is no longer visible (retro cue) (Souza and Oberauer 2016), suggesting it is the memory itself which is being manipulated and not simply attention to the sample.

One of the mechanisms by which the brain is proposed to facilitate this allocation of memory is through alpha oscillations; neural rhythms of around 10 Hz which are considered to be inhibitory in nature, supressing neural populations (Klimesch et al. 2007; Jensen and Mazaheri 2010). This theory posits that alpha oscillations may play an active role in the management of WM-related resources, inhibiting/activating those areas of the brain pertaining to irrelevant/relevant short-term memories. Indeed, electroencephalography (EEG) and magnetoencephalography (MEG) studies have shown that a directional cue (i.e., left or right) induces a retinotopic re-distribution of alpha power during WM retention (Sauseng et al. 2009; Poch et al. 2014; Foster et al. 2015; Myers et al. 2015; Leenders et al. 2018; van Ede et al. 2019a; Mössing and Busch 2020), i.e., in the case of a leftward cue, alpha power will be greater over the left visual cortex which is not needed (ipsilateral inhibition) than over the right visual cortex which is needed (contralateral activation) during WM retention.

Although this has been a remarkably popular WM paradigm, and the lateralisation of alpha power during cued WM retention has been replicated many times, a strong link between the physiology and the behaviour has not been established – in fact, in a thorough and well-powered investigation on the matter, Mössing and Busch found no association between the lateralisation of alpha power and any subsequent behavioural parameters (Mössing and Busch 2020).

Elsewhere the frequency of alpha oscillations has been more directly implicated in cognition. Alpha frequency increases as a function of cognitive effort (Klimesch et al. 1993; Ergenoglu et al. 2004; Haegens et al. 2014; Maurer et al. 2015; Mierau et al. 2017), correlates with WM performance (Richard Clark et al. 2004; Grandy et al. 2013; Puttaert et al. 2021), appears slower with ageing (Richard Clark et al. 2004; Mierau et al. 2017; Donoghue et al. 2020; Auer et al. 2024), and in certain pathologies, e.g. schizophrenia (Ramsay et al. 2021) and mild cognitive impairment (Garcés et al. 2013). However, the presence, or contribution, of frequency lateralisation in WM tasks has not been explored, nor have there been attempts to modulate alpha frequency during these tasks in a hemifield/hemisphere-specific manner. Whilst there have been some investigations of this sort on perception, i.e., by deploying brain stimulation approaches such as transcranial magnetic stimulation (TMS) (Di Gregorio et al. 2022) or transcranial alternating current stimulation (tACS) (Wöstmann et al. 2018; Deng et al. 2019; Battaglini et al. 2020) to modulate alpha frequency in the context of perception, alpha frequency is far less commonly the target in WM neuromodulation experiments.

Here, we sought to synthesise these two principles of alpha lateralisation and alpha frequency. Since alpha frequency is positively correlated with WM performance, and visual WM is thought to be facilitated by diminished alpha oscillations in the contralateral (and amplified alpha oscillations in the ipsilateral) visual cortex, we hypothesised that the rightward lateralisation of alpha frequency (i.e., faster over right hemisphere than left) during WM retention would be positively correlated with the leftward lateralisation of subsequent recall (i.e. left cues remembered better than right) – see **Figure 1**.

**Figure 1.**
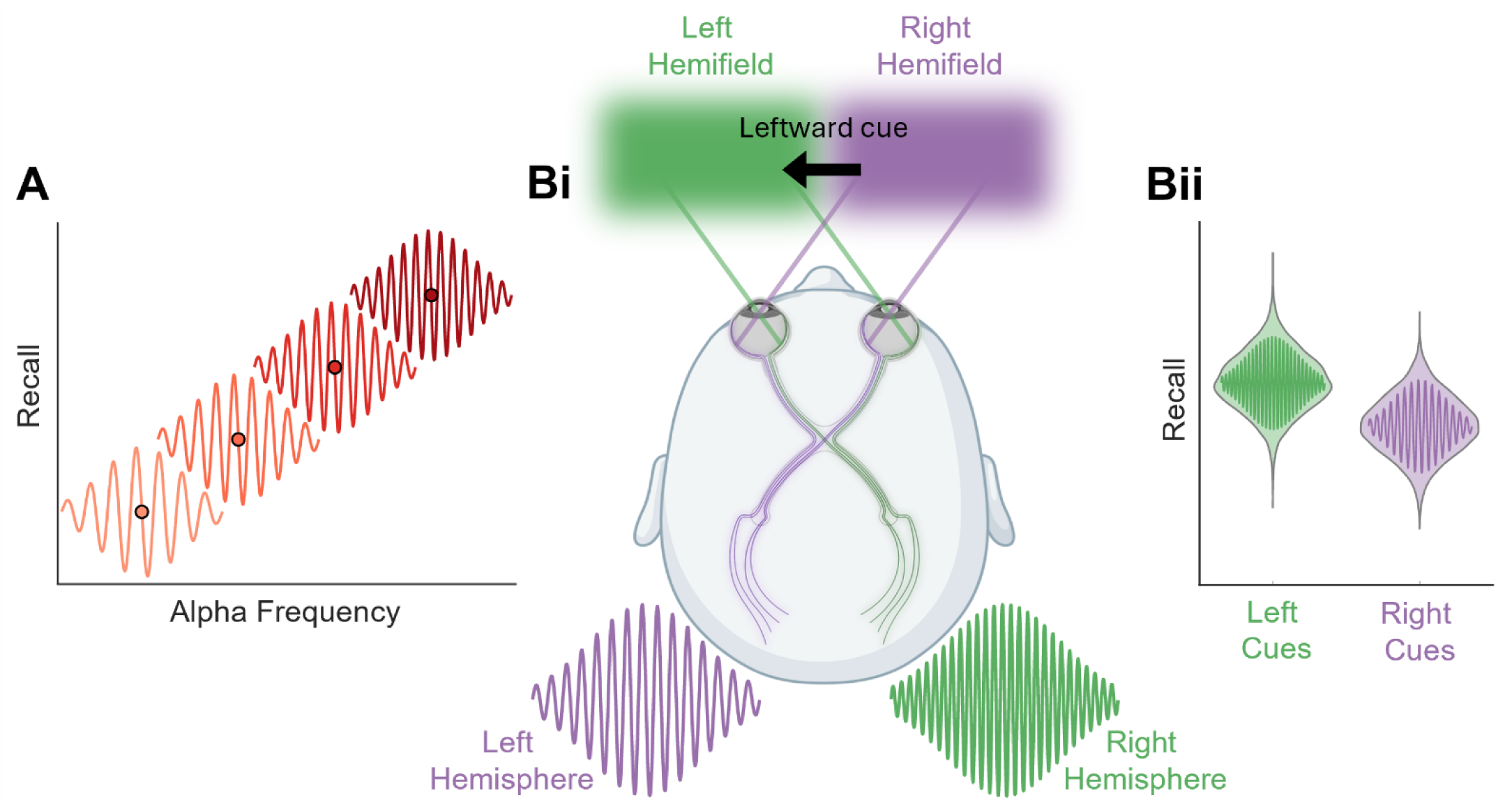
Visual representation of hypotheses **(A)** Alpha frequency is expected to correlate with memory performance **(B) (i)** Alpha frequency is expected to lateralise in a retinotopic manner, in this case a leftward cue is expected to increase frequency in the right (contralateral activation) hemisphere **(ii)** The extent of alpha frequency lateralisation is expected to correspond to the equivalent lateralisation in memory recall – i.e., faster right hemisphere alpha, better recall of left-cued memories.

To instigate and investigate this lateralisation of posterior alpha frequency, we employed alpha Closed-Loop Auditory Stimulation (αCLAS), which we have previously demonstrated to modulate alpha frequency in both wakefulness (Hebron et al. 2024) and REM sleep (Jaramillo et al. 2024) in a phase-dependent and spatially-specific manner. Here, we specifically sought to push apart the frequency of left and right hemispheres, so as to target opposite phases of alpha oscillations over opposite occipito-parietal hemispheres.

We hypothesised that:

(1) αCLAS can be used to target opposite phases over opposite hemispheres;
(2) this will result in phase-dependent alpha frequency changes, pushing the hemispheres apart in their frequency;
(3) task performance will be positively related to alpha frequency during the retention period;
(4) the lateralisation of alpha frequency (i.e., right hemisphere faster than left) during the retention period will be positively related to the lateralisation of behaviour (i.e. right-cued trials better remembered than left-cued);
(5) αCLAS will cause behavioural performance to be more lateralised than in the OFF condition, in accordance with hypothesis 4.

## 2. Materials and methods

### 2.1 Ethical statement

All experiments were approved by the University of Surrey Ethics Committee (approval numbers: FHMS 19–20 103EGA and FHMS 22–23 123EGA) and conducted in accordance with the Declaration of Helsinki. All participants gave written informed consent and were compensated for their time.

### 2.2 Experimental design

Two experiments were conducted, differing only in the participants involved, and the referencing scheme used for the real-time phase estimation. In experiment 1, electrode P4 was referenced to electrode P3 (**Figure S1A**), and in experiment 2 the opposite scheme was used (**Figure S1B**).

### 2.3 Participants

At the School of Psychology of the University of Surrey, 46 participants took part in this study, 21 in experiment 1 and 25 in experiment 2. One participant was removed from each experiment due to poor performance (< 50% accuracy). One participant was excluded from experiment 1 due to issues with the button box recording the responses. Four participants were removed from experiment 1 due to excessive noise in the EEG. One participant was removed from experiment 2 due to missing EEG markers. This left a sample of 19 participants in experiment 1 (12 female, mean ± std age = 24.411 ± 5.75 years) and 23 participants in experiment 2 (13 female, mean ± std age = 22.09 ± 2.15 years). Four participants overlapped between the two experiments.

### 2.4 Experimental protocol

Participants sat in a soundproof room with constant LED light (800 Lux), a chin rest was used to ensure constant distance (~60 cm) and orientation to the screen. Participants were exposed to 6 blocks (3 blocks per condition) of 50 trials of a visual working memory task based on (Leenders et al. 2018) (see **Figure 2).** Each trial consisted of a baseline (2 seconds), a cue (left or right arrowhead, 0.5 seconds), a sample (to memorise, 0.1 seconds), a retention period (5 seconds), and a probe (i.e, change or no change, 1 seconds), in that order. The sample to be remembered consisted of 6 coloured squares, 3 on each side of the screen. The ‘change’ involved one of the squares on the cued side changing colour, in the ‘no change’, the probe exactly matched the sample, the un-cued side never changed. These parameters were randomised in their order and counter-balanced such that each combination of left-cue, right-cue, change, and no-change made up around 25% of total trials. The task was coded in and presented using PsychToolbox (https://github.com/Psychtoolbox-3). Answers were given using a custom-made button-box, with two buttons. Participants pressed the left button to indicate that the probe matched the sample and the right that it did not. Participants were instructed to maintain fixation in the centre of the screen, so as to ensure the left hemifield was exposed only to the left side of the screen. Conditions differed only in the retention period, during which auditory stimulation was applied in the ON condition, whereas no sounds were played in the OFF condition. Conditions were alternated within each participant, e.g. [ON, OFF, ON, OFF, ON, OFF], and the starting condition was alternated between participants, e.g. participant 1: ON start, participant 2: OFF start, etc. Between blocks, electrode impedance was checked and, if necessary, adjusted. In total, each session lasted approximately 2 h.

**Figure 2.**
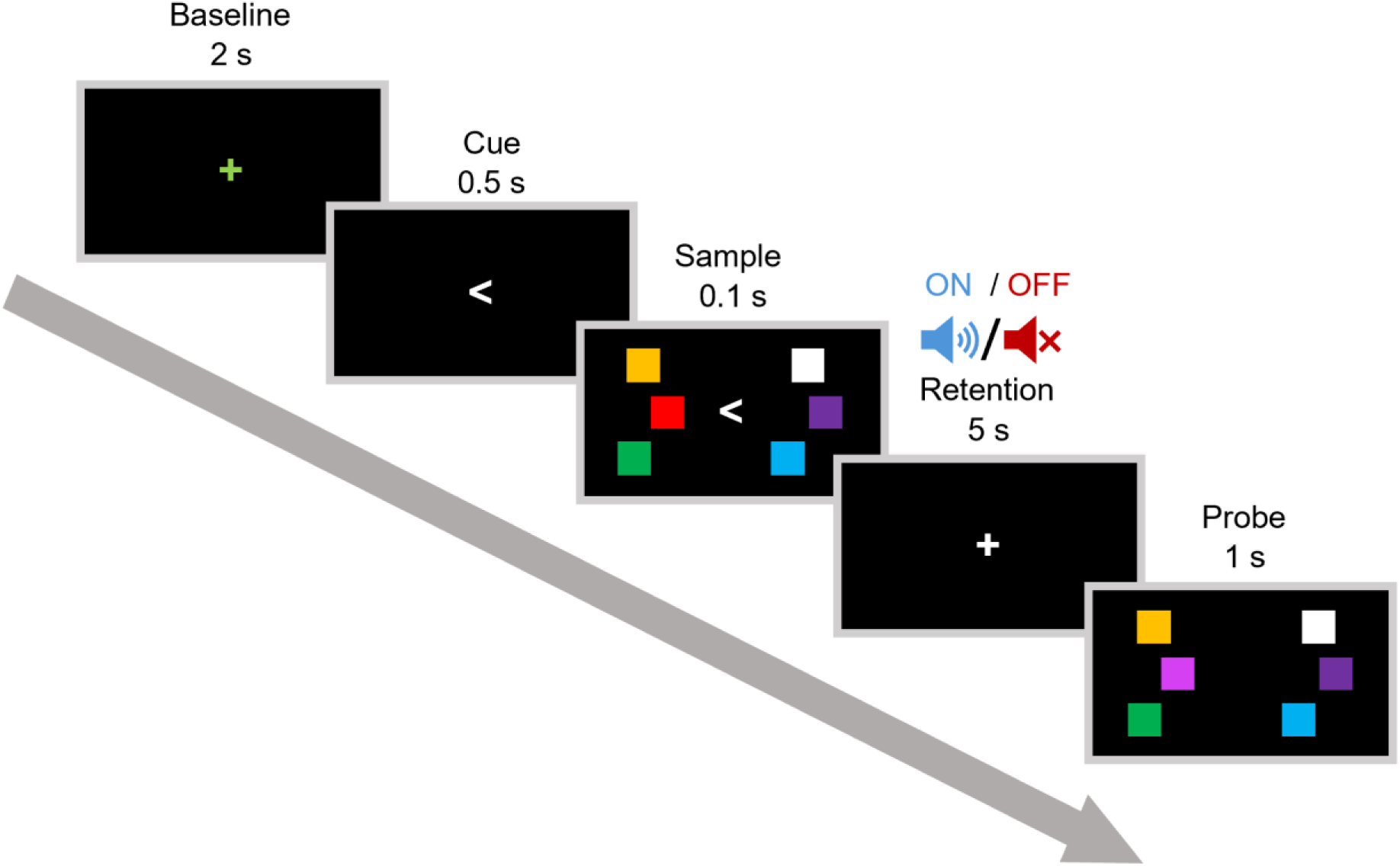
Task trial scheme. Each trial followed an identical format, cues can be left or right, probe can be identical or different from sample, this trial has a left cue and the sample changed. Conditions (ON and OFF) differed only during the retention period, during which stimulation was administered or not.

### 2.5 Phase-Locked Auditory Stimulation

Repetitive-αCLAS was applied as 20 ms pulses of pink noise, phase-locked to 7.5-12.5 Hz EEG rhythms, and delivered via in-ear etymotic headphones (Etymotic model ER-2). The endpoint-correct Hilbert Transform (Schreglmann et al. 2021) was used for accurate real-time phase-locking of EEG oscillations, as we have done previously (Hebron et al. 2024; Jaramillo et al. 2024). Sounds were targeted to 180° (i.e., the trough) in the signal electrode and a bipolar referencing scheme was employed (**Figure S1**), with the objective of applying sounds at opposite phases over opposite hemispheres. In experiment 1, electrode P4 was referenced to electrode P3 (**Figure S1A**), and in experiment 2 the opposite scheme was used (**Figure S1B**). Stimulation was applied exclusively during the retention period of the task, and the two experiments differed only in the referencing scheme and the specific participants taking part.

### 2.6 Behaviour Analysis

We computed d prime (Green and Swets 1966) (also, see supplementary materials, **Figure S2**), as a metric of memory performance which seeks to remedy response biases and offers a standardised score.

### 2.7 EEG/EOG data acquisition

High-density (hd)-EEG recordings were acquired in parallel with the phase-locking EEG device. Recordings were obtained using an actiChamp-Plus amplifier (Brain Products GmbH) and 128-channel water-based R-NET electrode cap (EasyCap GmbH, Brain Products GmbH) positioned according to the international 10-20 system. Electrode impedances were below 50 kΩ at the start of each recording, as per the manufacturer’s guideline, and were adjusted between each run, to ensure accurate impedances throughout the recording. All channels were recorded referenced to Cz, at a sampling rate of 500 Hz.

Additionally, horizontal electrooculography (hEOG) was employed to monitor, during data collection, that participants remained fixated on the centre of the screen. If participants moved their eyes, the block was restarted. EOG data were not subjected to post-hoc analysis.

### 2.8 EEG Pre-processing

All EEG data were pre-processed and analysed in MATLAB 2021a (Mathworks, Natick, MA). EEG filtering was carried out using the FieldTrip toolbox (Oostenveld et al. 2011), all other pre-processing was carried out using EEGLAB v2022.0 (Delorme and Makeig 2004). All topoplots and related statistics were produced using FieldTrip (Oostenveld et al. 2011).

Hd-EEG data were high-pass filtered (1 Hz) and notch filtered (50 Hz, 100 Hz), before automatic subspace reconstruction (ASR) was used to interpolate noisy channels. Artefacts were visually identified and removed. This was followed by independent component analysis, to identify and remove components related to ocular and cardiac artefacts.

### 2.9 EEG Analysis

Accuracy of phase-locking was calculated for each channel, participant, condition, and experiment using the following method. For every stimulus, the EEG was ‘epoched’ in 1 s windows up to and including stimulus onset, before the ecHT algorithm was applied offline, and the final phase estimate in each window was taken. The resultant vector length of these values was then computed, giving a value between 0 and 1, representing perfectly uniform and perfectly unimodal circular distributions respectively. The accuracy of phase-locking for the group was summarised by computing the average resultant of all participants, and computing a z-test. Z-tests were computed using average phase angle and resultant per participant, and the respective target phase angle (i.e., 0°, 180°).

For all analyses regarding power, the same method was used; power was computed in 1-second epochs, 2-17 Hz, in steps of 0.1 Hz, Hamming window, using the ‘pwelch.m’ MATLAB function. For all analyses regarding frequency, Cohen’s instantaneous frequency sliding method was used (Cohen 2014).

In order to reduce the dimensionality of the data and to facilitate investigations into the interaction of stimulation, frequency, power, time, and performance, two electrode *clusters-of-interest* (COI) were identified. These were defined on the basis of which electrodes saw a significant change in alpha frequency as a result of sound stimulation. Since the only difference between experiments 1 and 2 was the hemispheres at which phase-locking was 0° or 180° (experiment 1: left 0°, right 180°; experiment 2: left 180°, right 0°), the entire topography of experiment 2 was mirrored, such that the left hemisphere and right hemisphere were inverted, and the experiments could be directly compared. Following this inversion, a cluster of channels on the right hemisphere (now 180° in both experiments) was found to have a significantly higher alpha frequency than OFF. This cluster was labelled 180COI (180° Cluster of Interest) and the equivalent electrodes on the opposite hemisphere were taken as 0COI (0° Cluster of Interest). Additionally, there are two instances in which we were interested in the genuine topography for use in correlations (**Figures 6** and **7**), i.e., without flipping experiment 2. These right and left clusters are referred to as right cluster-of-interest (RCOI) and left cluster-of-interest (LCOI), respectively. These clusters are frequently used in the analysis.

### 2.10 Statistical Analysis

All statistics are described in the figure legends, but briefly: in cases of topographical plots, cluster-corrected t-tests or correlations were used, except in the case of phase-locking accuracy in which Bonferroni correction was employed. Otherwise, t-tests (paired, two sample, and one sample) and simple linear regression make up the remainder of the tests employed.

## 3. Results

### 3.1 Sounds were accurately phase-locked to opposite phases of alpha on opposite hemispheres

In experiment 1, sounds were accurately locked to 0° over the left parietal/occipital hemisphere, and to the opposite phase over the equivalent area of the right hemisphere, as assessed by z-test (**Fig 3Ai**). In experiment 2, with an inverted referencing scheme, the opposite phase relationship was observed between hemispheres (**Figure 3Aii**). Phase angles and resultants were highly consistent across participants and experiments (**Figure 3B**). A clear dynamic in the mean inter-stimulus interval (ISI) of stimuli was observed across the retention period for each trial, such that frequency increased over the first ~10 stimuli, before gradually dropping to below its starting ISI by the 40^th^ stimulus (**Figure 3C**). This dynamic was consistent across ON and OFF (in which markers were logged when a stimulus would have been administered, but volume was 0 dB) conditions, with a slightly higher mean ISI in experiment 1 than in experiment 2. ISIs were stable across *trials* within a block, indicating that the frequency at which the sound was delivered (or logged for the OFF condition) was stable over time, (**Figure 3D**). ISI and alpha frequency were strongly related in both conditions and experiments, such that approximately 1 stimulus was administered per alpha cycle.

**Figure 3.**
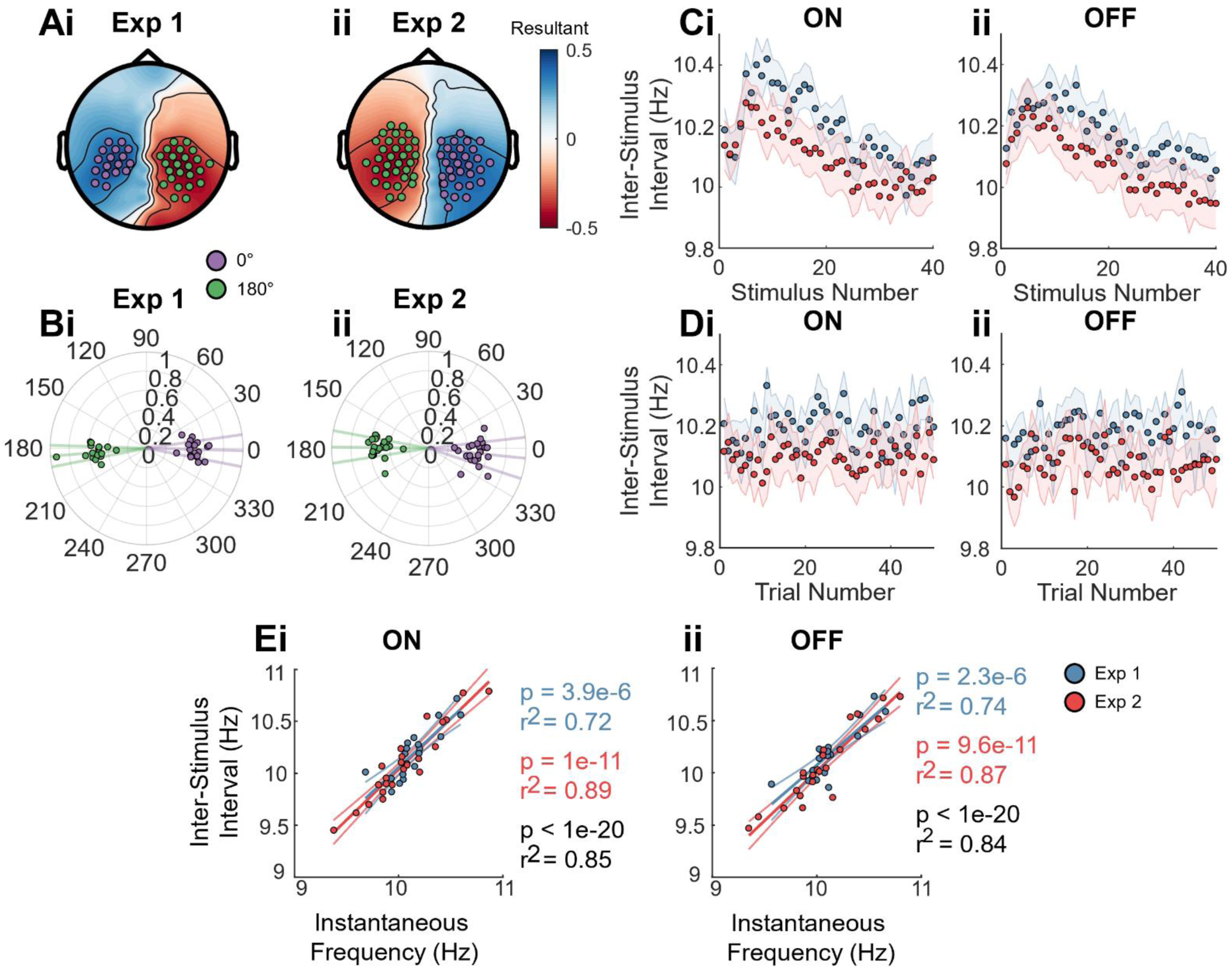
Characterisation of stimulation. **(A)** Topography of phase-locking resultant at stimulus onset in **(i)** experiment 1 and **(ii)** experiment 2. Note: the sign of resultant is reversed for each channel at which the average phase angle is closer to 0° than 180°. Purple circles represent channels at which stimuli were significantly unimodal around 180°, and green circles show the same for 0°, as per Bonferroni-corrected z-test. **(B)** Polar plots showing the phase angle and resultant for each participant in **(i)** experiment 1 and **(ii)** experiment 2. Data were taken from the channels identified as significantly unimodal in **1A** for 180° (purple) and 0° (green) and averaged within each participant. Scattered points indicate individual participants, lines indicate the mean phases angle and the standard error of the mean. **(C)** The inter-stimulus interval (ISI) (i.e. frequency of stimulation) across the first 40 stimuli in the retention period for experiment 1 (blue) and experiment 2 (red), shown for both **(i)** ON and **(ii)** OFF conditions (in OFF, markers were logged for each sound stimulus, but volume was set to zero). Shaded error bar indicates standard error of the mean **(D)** Same as **1C**, but across the 50 trials of the block, averaged across blocks, rather than across the individual sound stimuli **(E)** Instantaneous alpha frequency vs ISI for both experiment 1 (blue) and experiment 2 (red), shown for **(i)** ON and **(ii)** OFF conditions. Alpha frequency is the mean frequency across all channels in 0 and 180 clusters-of-interest (see *Methods*). Scattered points indicate individual participant, fit line and statistics are taken linear regression of instantaneous frequency vs ISI which was run for experiment 1 (blue), experiment 2 (red), and both combined (black).

### 3.2 Alpha frequency increased during the retention of memory and correlated with performance

Before assessing the impact of sound stimulation on alpha frequency, we considered it important to first describe its natural trajectory and relationship to behaviour in the OFF condition, to establish a meaningful frame of reference.

Consistent with the dynamics of the inter-stimulus interval, instantaneous alpha frequency increased across the scalp at the beginning of the retention period, before falling to a lower level by second 5, relative to baseline (**Figure 4A**), indicating, that the stimulator had followed the brain. This frequency increase was correlated with d-prime (**Figure 4B**), most strongly in seconds 2 and 3 of the retention period, meaning those participants whose alpha frequency increased more performed better in the memory task. Frequency measured in central/motor areas appeared to reflect a separate process to that of the frontal/occipital alpha, which was correlated with performance. Whilst increases in alpha frequency emerged first over the central/motor areas (**Figure 4Ai**), the increase in these areas did not correlate with performance, unlike almost all other regions (**Figure 4B**).

**Figure 4.**
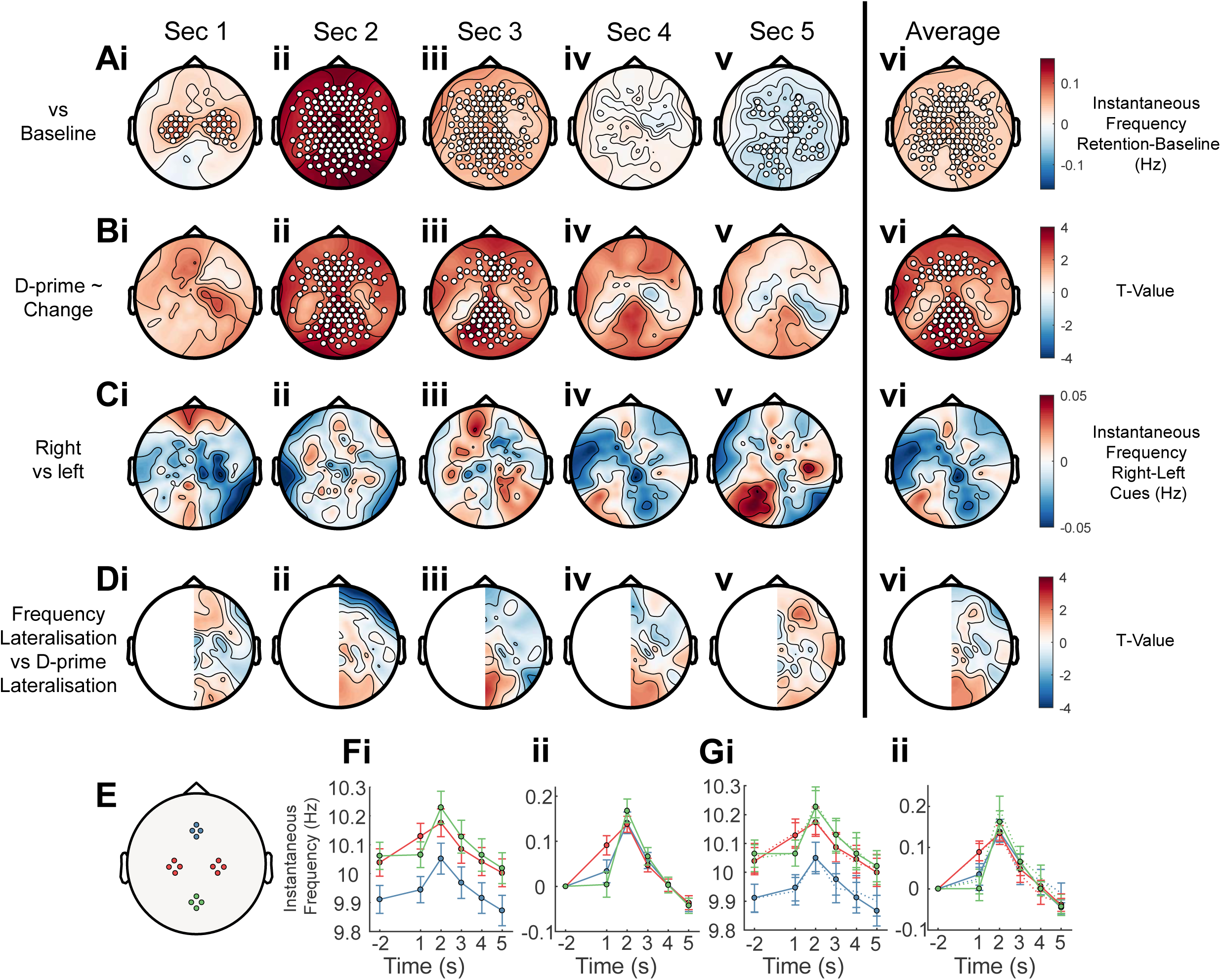
Characterisation of instantaneous alpha frequency in OFF condition. **(A)** Topographical plots for frequency differences between baseline (−2.5 to −0.5 seconds) and seconds 1 to 5 **(i-v)** of the retention period, as well as the average of all five seconds **(vi)**. Data are combined across experiments 1 and 2, with participants common to both experiments averaged across experiments, N = 38. Scattered points indicate cluster-corrected significant differences (*p*<0.05) from baseline, as calculated by permutation paired t-test. **(B)** Same as **5A**, but here permutation *correlations* were run between frequency change and d-prime, for each second of the retention period **(i-v)** and for their average **(vi)**. **(C)** Topographical plots for frequency differences between left and right-cued correct trials, for each second of the retention period **(i-v)** and their average **(vi).** No significant differences were observed by cluster-corrected permutation paired t-test. **(D)** Topographical plots for correlation of frequency lateralisation (right hemisphere channels minus left hemisphere) and d-prime lateralisation (left-cued trials minus right cued-trials; all trials correct and incorrect). No significant correlations were observed **(E)** Schematic of four clusters of electrodes, chosen to represent four different brain regions: frontal (blue), central/motor (red), and occipital (green) **(F)** Time-series of instantaneous alpha frequency for each cluster, following the aforementioned colour scheme during the retention period. Data are shown for **(i)** absolute frequency and **(ii)** baseline-subtracted frequency. **(G)** Same as **4A** but for correct (solid lines) and incorrect (dashed lines) trials.

No difference in alpha frequency was observed when contrasting correct left-cued and right-cued trials (**Figure 4C**), nor was there a correlation observed between the lateralisation of frequency (right hemisphere – left hemisphere)) and the lateralisation of d-prime (right cues minus left cues) (**Figure 4D**). This suggests that alpha frequency does not ordinarily play a prominent retinotopic role in WM.

Time series of alpha frequency from frontal, central/motor, and occipital regions (**Figure 4E**) showed slightly different trajectories. Frontal alpha clearly had a lower frequency throughout the retention period, and as outlined above, central alpha increased most rapidly (**Figure 4Fi**). Interestingly, when each region was normalised to the baseline (baseline-subtracted), a near identical trajectory was observed in seconds 2-5 of the retention period, meaning a highly consistent *relative* change in alpha frequency was conserved across the scalp (**Figure 4Fii**).

Plotting the same time series following segregation of correct and incorrect trials revealed next to no difference between trials that were remembered and those that were not (**Figure 4G**), suggesting that the group-level correlations in **Figure 4B** were reflective of a trait-like individual difference, rather than trial-by-trial differences.

### 3.3 Sounds altered alpha frequency in a phase-dependent and spatially localised manner

Next, we wanted to see if alpha frequency was modified according to our model mechanism of αCLAS.

Indeed, consistent with our previous work (Hebron et al. 2024; Jaramillo et al. 2024), αCLAS modulated frequency in a spatially-specific and phase-dependent manner, relative to OFF. In both experiments, the hemisphere over which sounds were phase-locked to 180° saw a small but consistent increase in frequency (**Figures 7Ai**, **5Ai**, **5Aii**). Since the phase-locking accuracy and behavioural performance did not differ between experiments 1 and 2 (see **Figure 8**), they were combined, more clearly demonstrating this effect (**Figure 5Aiii**). Sounds locked to 0° (contralateral hemisphere), meanwhile, did not appear to modulate frequency.

**Figure 5.**
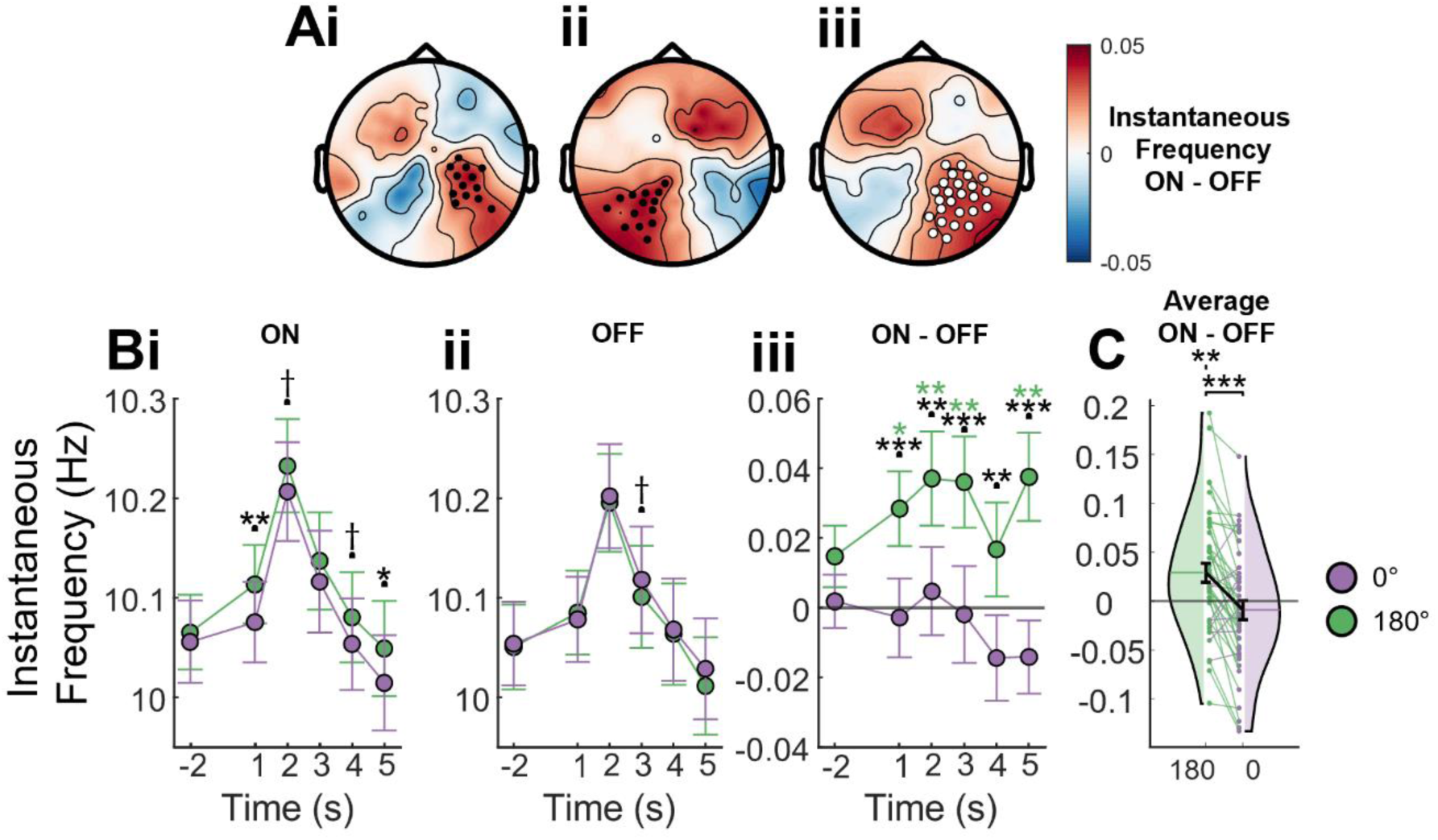
Stimulation-induced changes to alpha frequency. **(A)** Topographical plots of differences between ON and OFF conditions, averaged across the retention period, for **(i)** experiment 1, **(ii)** experiment 2, and **(iii)** Both experiments combined (note: when combining experiments, all channels are inverted across the midline in the case of experiment 2, since it is the mirror image of experiment 1, see *materials and methods*). Scattered points show cluster-corrected significant differences, as computed by permutation paired t-test (black .05>*p*<.1, white *p*<.05). The cluster seen here is used in several analysis, as is its mirror opposite, and they are referred to as 180° cluster-of-interest **(180COI)** and 0° cluster-of-interest **(0COI)** respectively. **(B)** Time series of absolute alpha frequency (mean and SEM) of 180° (green) and the 0° (purple) across the retention period, for **(i)** ON and **(ii)** OFF conditions. **(iii)** shows the ON-minus-OFF time series. Black statistics marks indicate result of paired t-tests between 180° and 0°. Green statistics marks indicate results of one-sample t-test (i.e., contrast with OFF) for 180°, and purple marks show the same for 0° **(C)** OFF-subtracted frequency, averaged over the full retention period. Violins show distribution of participants, scattered points show individual participants, lines show within participants, between conditions differences, error bars indicate standard error of the mean. Statistics bar between violins indicate significant paired t-test, statistics marks over violins indicate significant one-sample t-test, i.e., difference from OFF.

Time-series plots showed that the electrode cluster identified in **Figure 5Aiii** (180° cluster-of-interest (180COI), see M*ethods*) and its opposite (0° cluster-of-interest (0COI)) followed the same course as that identified in **Figure 4F**, in both ON (**Figure 5Bi**) and OFF (**Figure 5Bii**) conditions. Differences between clusters were observed in the ON but not the OFF condition throughout the retention period in the 180COI, but not the 0COI (**Figure 5Biii**). When averaging across the retention period it was observed that 29 of the 38 participants (76%) showed a higher alpha frequency in the 180COI than in the 0COI, relative to OFF (**Figure 5C**).

### 3.4 There was no main effect of stimulation on task performance

It was important to check that sound stimulation did not have a net negative effect on task performance, since such stimuli may serve as a distraction. Moreover, we had hypothesised that a lateralisation of alpha frequency might result in a lateralisation of behaviour. Memory performance, as measured by d-prime, did not differ between ON and OFF conditions (**Figure 6A**). The lateralisation of d-prime (i.e. right-cued trials minus left-cued trials) was computed to explore whether the group had an innate preference for left or right trials and whether stimulation modulated this, but again no significant effect was found.

**Figure 6.**
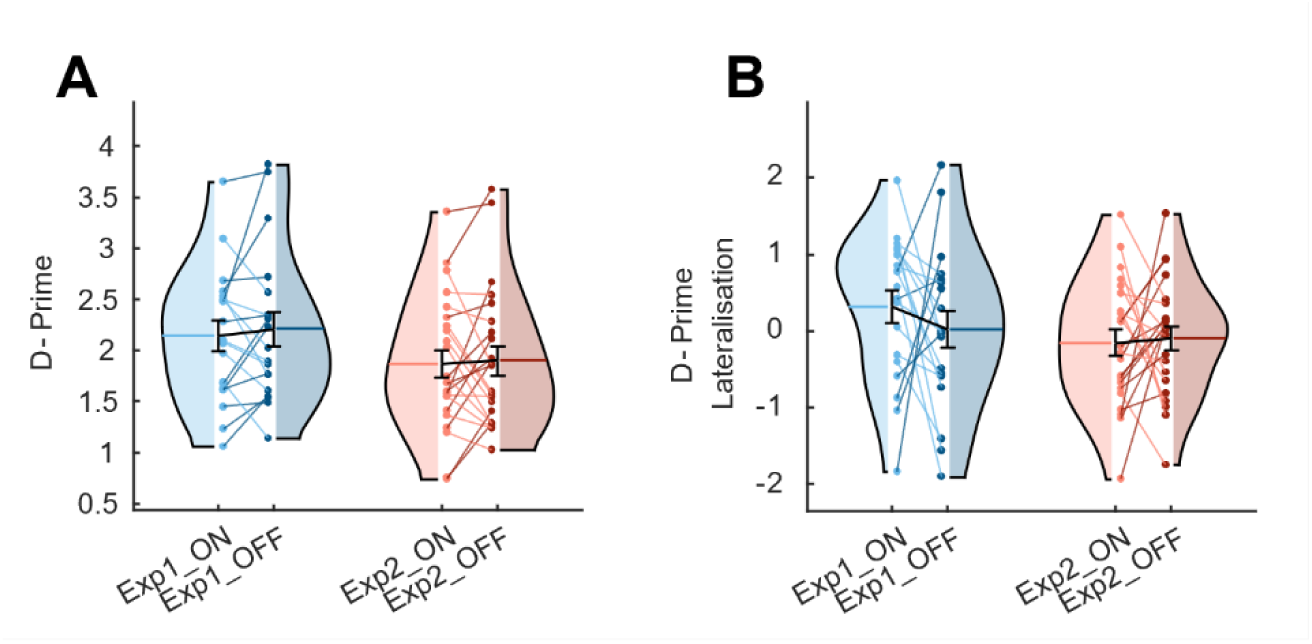
Task Performance. **(A)** Average d-prime across conditions and experiments. **(B)** The same as **A**, but for the lateralisation of d-prime, i.e. d-prime in right-cued trials minus d-prime in left-cued trials. Paired T-tests were used to compare conditions (ON/OFF) but no significant differences were observed Colour indicates the experiment and condition: ON in experiment 1 (bright blue), OFF in experiment 1 (dull blue), ON in experiment 2 (bright red), and OFF in experiment 2 (dull red). Violins show distribution of participants, scattered points show individual participants, lines show within participants, between conditions differences, error bars indicate standard error of the mean.

### 3.5 Degree of stimulation-induced frequency lateralisation correlated with lateralisation of performance and was dependent on the power of alpha oscillations

Although there was no main effect of stimulation on the lateralisation of behaviour, we considered that this may result from the heterogeneity of effects of stimulation on frequency – whilst it was clear that 180-αCLAS induced a faster alpha frequency than 0-αCLAS, the extent to which this occurred differed between participants (**Figure 5C**). Hence, we enquired whether those individuals for whom the physiological effect of stimulation was larger also saw a greater behavioural effect.

Indeed, the stimulation-induced lateralisation of frequency (right hemisphere – left hemisphere) was positively correlated with that of d-prime (right cues – left cues), but only in second 3 of the retention period (**Figure 7A**). This relationship held for both experiments (**Figure 7B**) and, accordingly, when participants were sorted on the basis of their frequency lateralisation in second 3, a significant difference in d-prime lateralisation was observed between groups (**Figure 7C**) – such that those participants whose alpha frequency was more left-lateralised by stimulation (group 1) also saw their performance more left-lateralised, and vice versa. When performance was sorted in this fashion across time, both the correlation and the difference were statistically significant for much of the retention period (**Figure 7D**).

**Figure 7.**
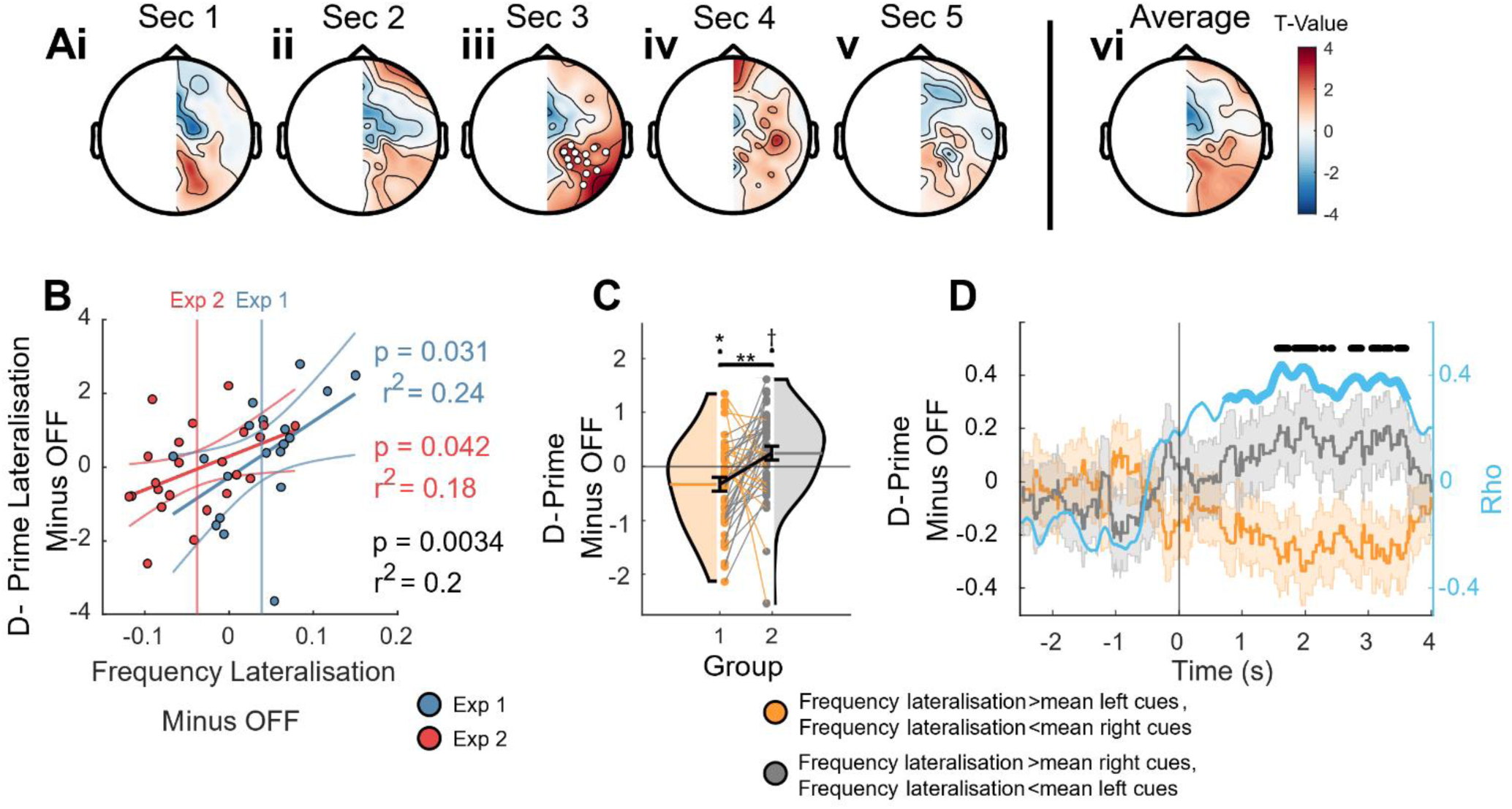
Relationship between lateral differential of alpha frequency and lateral differential of task performance. **(A)** Topographical plot of frequency lateralisation (right hemisphere minus left hemisphere) and d-prime lateralisation (right cues minus left cues) for seconds 1-5 **(i-v)**, and the average **(vi)**, of the retention period, for both experiments combined. Note: in this case, the topography of experiment 2 is not inverted across the midline before combination of experiments, since we wanted to investigate the actual left vs right lateralisation on the equivalent in the behaviour and each experiment pushed the brain in opposite directions. Scattered points show cluster-corrected significant (*p* <.05) relationships, as computed by permutation paired correlation **(B)** Scatter plot and linear fits of stimulation induced frequency lateralisation at the cluster(s)-of-interest ((stimRCOI – stimLCOI) - (OFFRCOI – OFFLCOI)) vs stimulation induced d-prime lateralisation ((stim_right_cues – stim_left_cues) - (OFF_right_cues – OFF_left_cues)) for second 3 of the retention period, for experiment 1 (blue) and experiment 2 (red). Scattered points indicate individual participants, vertical lines indicate average frequency lateralisation for experiments 1 and 2, blue and red respectively. Statistics show results of linear regression for experiments 1 (blue), experiment 2 (red), and both combined (black). **(C)** The stimulation-induced d-prime values (left- and right-cued trials, relative to OFF) of participants were sorted on the basis of their stimulation-induced frequency lateralisation (right-left) in second 3. Those whose frequency was more right-lateralised than the mean had their left cues sorted into group 1 (orange) and their right cues sorted into group 2 (grey), the opposite was carried out for participants whose frequency was less right-lateralised than the mean. Violins show distribution of participants, scattered points show individual participants, lines show within participants, between conditions differences, error bars indicate standard error of the mean. **(D)** The equivalent of **C**, but over time, i.e. rather than taking second 3 isolation, groups 1 and 2 are defined on the basis of a moving 1 second window, across the retention period. Black marks indicate significant paired t-test (*p* < .05) and the blue line shows rho value from correlation computed at each time point.

Interestingly, this relationship was in the opposite direction to our predictions; we had hypothesised that memory would be improved if frequency was higher contralateral to the cue.

We were also curious about the potential determinants of an individual’s amenability to αCLAS and, based on our previous findings (Hebron et al. 2024; Jaramillo et al. 2024) and oscillator theory (Winfree 2001), we proposed that it might depend on the magnitude of the oscillations themselves.

Indeed, the extent to which stimulation lateralised frequency was dependent on the endogenous individual alpha power (**Figure 8A**), as measured during the retention period in the OFF condition, in addition to the phase at which sounds were applied. This relationship between power and frequency change was found to be specific to power in the alpha band (**Figure 8B**) but did not show marked spatial specificity (**Figure 8C**).

**Figure 8.**
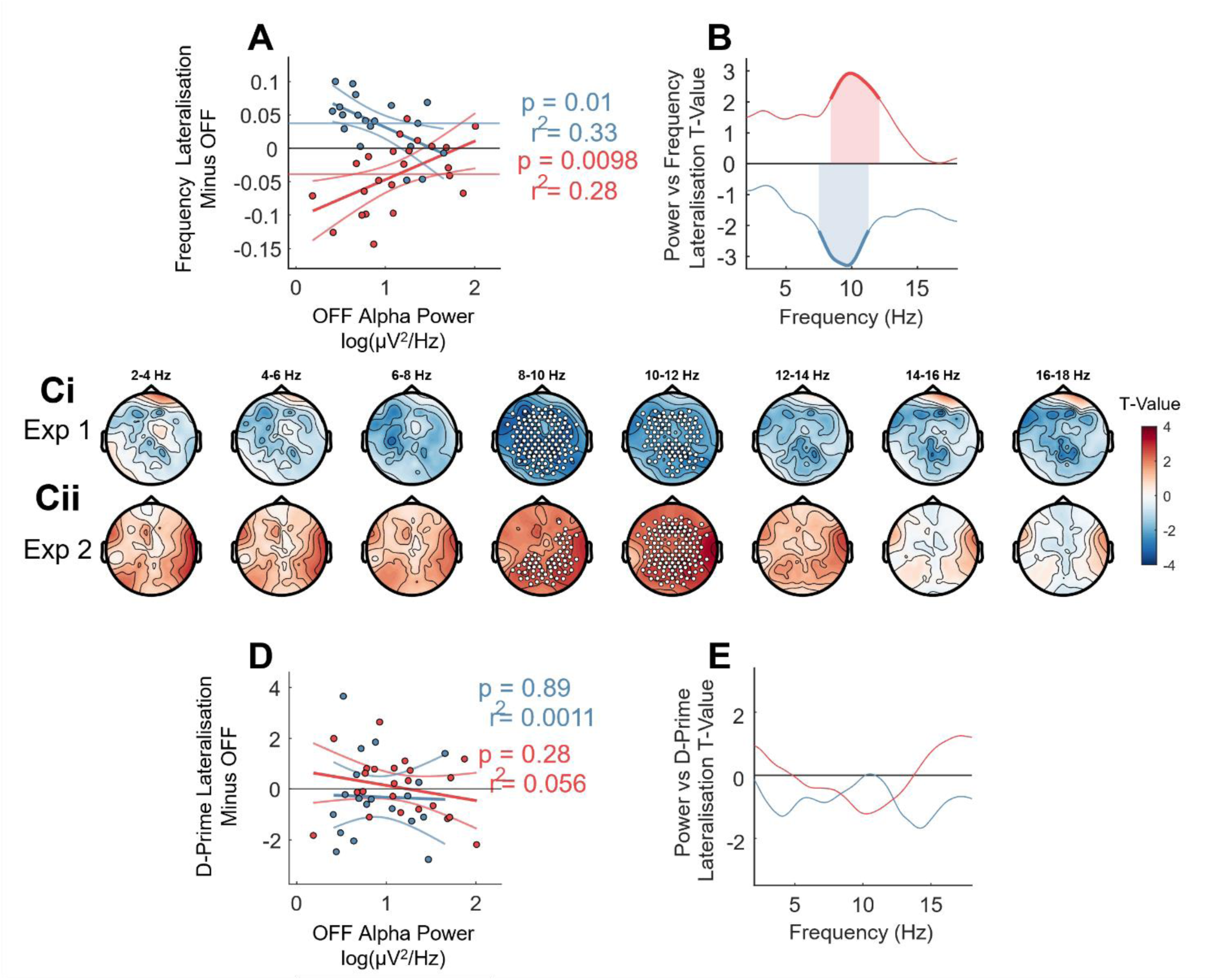
Relationship between alpha power, lateral differential of alpha frequency, and lateral differential of task performance. **(A)** Alpha (8-12 Hz) bandpower (log) taken from OFF, averaged across the retention period, and averaged across both the RCOI and the LCOI, vs stimulation-induced alpha frequency lateralisation ((stimRCOI – stimLCOI) - (OFFRCOI – OFFLCOI)). Scattered points indicate individual participants, fitted lines and statistics derived from linear regression for experiment 1 (blue) and experiment 2 (red), horizontal lines indicate average frequency lateralisation for experiment 1 (blue) and experiment 2 (red) **(B)** Same as linear regression from **A**, but instead of taking the full alpha band (8-12 Hz), regressions were run for each frequency, 2-17 Hz, in steps of 0.1 Hz. Bold, shaded portions indicate where those regressions were statistically significant in experiment 1 (blue) and experiment 2 (red). Horizontal line shows zero. **(C)** topographical plots of correlations between power and alpha frequency lateralisation, both averaged across the retention period, for **(i)** experiment 1 and **(ii)** experiment 2. Scattered points indicate cluster-corrected significant correlations (*p* < .05). **(D) (E)** Same as **A** and **B** but power is regressed against the stimulation-induced lateralisation of d-prime.

We supposed that the endogenous alpha power of an individual might then mediate the effect of stimulation on behaviour. However, despite the correlation between frequency lateralisation and d-prime lateralisation, and the correlation between alpha power and frequency lateralisation, there was no such correlation found between alpha power and d-prime lateralisation (**Figure E8D**, **8E**).

## 4 Discussion

Here, we have shown: (1) that αCLAS can be used to phase-lock sounds to opposite phases of alpha on opposite hemispheres; (2) during WM retention in a classical alpha-related visual WM task, alpha frequency increases and the extent of this increase correlates with task performance; (3) αCLAS modulates posterior alpha frequency in a spatially-specific and phase-dependent way; (4) αCLAS-induced frequency modulation correlates with the lateralisation of performance; (5) the extent of frequency modulation is dependent on alpha power.

This represents several advances on our previous investigations (Hebron et al. 2024; Jaramillo et al. 2024), the first of which is the implementation of a bipolar-referenced phase-locking electrode to target opposite phases. The importance of choice of reference is well recognised (Cohen 2015), and here we use it to our advantage to improve the specificity of our neuromodulation approach. We have not observed this particular approach to closed-loop neuromodulation in the literature, although sometimes an online Laplacian is used in state-dependent TMS (Zrenner et al. 2018; Bergmann et al. 2019; Schaworonkow et al. 2019). Another advance is the possibility of modulating parietal and occipital alpha oscillations, which we were previously unable to do with repetitive-αCLAS. Sounds timed to 180° (i.e., the trough) increased the frequency of alpha rhythms, whilst on the opposite hemisphere (0°, peak) there were no frequency changes observed, which we suspect is because the peak represented the so-called ‘dead zone’ of the phase response curve of this particular oscillation, i.e. that phase at which a stimulus neither advances nor delays the oscillation (Johnson 1999). We had hypothesised that, previously, the inability of sound to interact with these posterior rhythms had less to do with anatomy (i.e. more visually specialised than auditory) and more to do with their amplitude. This was because oscillator theory suggests that perturbation of an oscillation is dependent on both the magnitude of the input and that of the oscillation (Winfree 2001), and alpha amplitude is greater over posterior regions. Here, the amplitude of alpha rhythms is lowered since, in contrast to our previous experiment (Hebron et al. 2024), the eyes are open, and we believe this underpins their successful modulation. This was also reflected at the individual-level, whereby participants with lower endogenous alpha power were more amenable to modulation (i.e. changes in alpha frequency) by αCLAS. These two points have important implications. One of which is that posterior alpha is far and away the most common target of alpha-focussed neuromodulation, and the ability to interact with oscillations in this region greatly boosts the potential use-cases of αCLAS. he second, potentially profound, implication is that an individual’s sensitivity to stimuli more generally may be determined by the amplitude of their oscillations.

The αCLAS-induced frequency changes themselves were comparable in magnitude to those effects reported as meaningful elsewhere (Cohen 2014; Samaha and Postle 2015; Wutz et al. 2018). They were, however, slight in their magnitude; ~ 0.05 Hz, compared to what we observed in our previous application of aCLAS (.05 Hz, Hebron et al. 2024; Jaramillo et al. 2024).

Whilst we previously focussed on the alpha rhythms of the resting or sleeping brain, here we target alpha during a process in which it is thought to play an active role. It appears that the same principles apply to αCLAS of these ‘active’ rhythms as their passive, or resting, equivalents. This is fundamentally important, if αCLAS is to be a useful tool for *cognitive* neuroscientists – for example, if one wants to either modulate alpha-related behaviour, assess the contributions of alpha to behaviour, or both.

With respect to the *neuroscience* questions at hand, several findings of note should be discussed. Firstly, there was a clear dynamic in the frequency of alpha oscillations across the retention of WM, a sharp increase followed by a gradual decline. This particular trajectory has, to our knowledge, not been reported before, most likely because we use longer retention periods than typically seen in the literature (i.e. 5 seconds, as opposed to ~1 seconds in most cases (Griffin and Nobre 2003; Foster et al. 2015; Myers et al. 2015; Wutz et al. 2018; Mössing and Busch 2020) and 2.5 seconds in one study (van Ede et al. 2019b)), which can capture this dynamic, and indeed because most studies neglect to investigate the frequency of pertinent oscillations. Importantly, alpha frequency was associated with behaviour. In agreement with the literature and our expectations, a greater increase in alpha frequency positively associated with WM performance on a between-participants basis, i.e. group correlation. However, correct and incorrect trials could not be distinguished by alpha frequency, within-participants. We interpreted this to mean that alpha frequency in OFF reflected a trait-like quality, which may relate to attention, vigilance, or indeed memory. Theoretical and computational accounts of this frequency shift suggest that alpha oscillations are generated by recurrently connected networks of spiking neurons and an *increase* in spiking activity leads to an *increase* in alpha frequency (Cohen 2014; Lefebvre et al. 2015; Mierau et al. 2017), which may explain our observations – crudely put, when there are more microscale computations occurring the macroscale oscillation appears faster.

Whilst alpha frequency positively correlated with recall generally, with a fairly global topography, the retinotopic effect in which we were most interested was not present in OFF. This suggested the spatial distribution of alpha frequency might not index that of the visual information being maintained in WM. However, the absence of an effect of αCLAS on behaviour might instead be explained by the minimal endogenous lateralisation of alpha frequency (i.e. a very narrow, near-zero distribution, both hemispheres are naturally very similar to one another) since the αCLAS-induced changes to frequency lateralisation did indeed correlate with that of behaviour.

Interestingly, this correlation was in the opposite direction to our expectations. We had hypothesised that (given frequency correlates with performance (Richard Clark et al. 2004; Grandy et al. 2013; Puttaert et al. 2021), replicated here) and visual WM is thought to be managed by the contra-lateral visual cortex, increasing the frequency over the right hemisphere would improve performance in left-cued trials and *vice versa*, but the opposite effect was observed. We suggest that 180° αCLAS may be well-timed to perturb alpha oscillations (hence the phase-specific frequency changes), but also to *disrupt* the associated processes, be they attention, memory, or perceptual sensitivity. Bonnefond and Jensen found that alpha resets to a particularly insensitive phase in anticipation of, and to protect against, predictable visual distractors (Bonnefond and Jensen 2012) – here we may be facilitating the precise opposite; timing stimuli to a particularly sensitive alpha phase (180°), so as to effectively “distract” ongoing processes over that hemisphere. If this is indeed the case, disentangling the contribution of frequency to behaviour and phase to cortical excitability using αCLAS could prove remarkably difficult, since it may leverage the latter to influence the former – frequency and behaviour change could both be consequent of a third, separate, upstream process. For this reason, we must remain cautiously agnostic to the role of alpha frequency lateralisation in the remembering of lateral visual information, for now.

There are two standout points of contention in this work. The absence of a group-level effect of αCLAS phase on behaviour and the absence of a relationship between alpha power and the behavioural effects of αCLAS. Since αCLAS modulates alpha frequency in a lateralised fashion, the extent of this frequency lateralisation correlates with the lateralisation of behaviour, and alpha power predicts the individual amenability to αCLAS modulation, one would expect behaviour to be modulated by αCLAS in a phase-dependent manner and for power to correlate with this behavioural change, but this was not the case. This discrepancy remains mysterious, but may arise from the relatively small effect of frequency on behaviour – if this effect was considerable one would expect it to be present as a main effect of stimulation on behaviour, which it was not. It may be the case that a larger sample, or a further optimised stimulation approach (e.g. real-time Laplacian reference (Zrenner et al. 2018; Bergmann et al. 2019; Schaworonkow et al. 2019) or personalised spatial filters (Schaworonkow et al. 2018) may be able to bring this effect to light.

Nevertheless, this study has, at the very least, opened up new questions regarding alpha oscillations and WM, and yielded many important insights into the use of αCLAS for neuromodulation.

## CRediT

**HH**: Conceptualization, Data curation, Formal analysis, Investigation, Methodology, Project administration, Software, Resources, Supervision, Visualization, Writing – original draft.

**RD**: Conceptualization, Data curation, Investigation, Writing – review & editing

**VJ**: Methodology, Writing – review & editing

**DJD**: Methodology, Supervision, Writing – review & editing

**IRV**: Conceptualization, Funding acquisition, Methodology, Project administration, Software, Resources, Supervision, Writing – review & editing

