## Supplementary materials for "Lateralised modulation of posterior alpha oscillations by closed loop auditory stimulation during memory retention"

### Referencing Montage for Closed-Loop

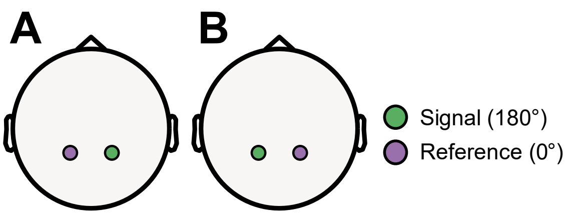


**Figure S1. Bipolar referencing montage.** Opposite referencing schemes were used for αCLAS in **(A)** experiment 1 and **(B)** experiment 2.

### D Prime

D prime is a measure of task performance which seeks to remedy response biases, i.e., if a participant only ever answers 'change', they will score 50% accuracy despite not engaging with the task, and hence percentage accuracy may not be representative of actual performance. D prime, on the other hand, is derived from hit rate (H) (proportion of 'change' answers when there is a genuine change) and false alarm rate (FA) (proportion of 'change' answers when there is no change). A normal distribution of potential proportions is assumed in both cases and each is converted into a z-score, the difference between these z-scores gives d prime (equation 1) – the larger its magnitude the better the performance. **Figure S2** shows the relationship between hit and false alarm rates and d prime.

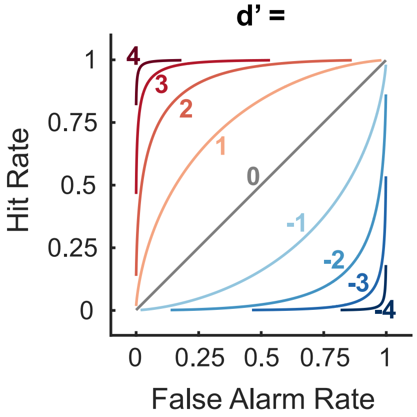


**Figure S2. d prime.** The various possible combinations and contributions of hit and false alarm rate that can lead to various (-4 to 4) values of d-prime. Lines indicate thresholds for values of d prime.

$$\boldsymbol{d' = z(H) - z(FA)}$$

**Equation 1.** Calculation of d-prime.
